# OsbHLH089 and OsbHLH094 Modulate OsSLR1 Levels to Maintain Male Reproductive Fitness in Rice

**DOI:** 10.1101/2024.11.24.624999

**Authors:** Telma Fernandes, Pedro M. Barros, María Flores-Tornero, Pedro Carvalho, Helena Sapeta, Jörg D. Becker, Isabel A. Abreu

## Abstract

DELLA proteins are a unique class of transcriptional regulators in plants, playing critical roles in diverse biological processes. Far from being only negative regulators of the Gibberellin (GA) signalling pathway, DELLAs act as central signalling hubs due to their versatile binding capacity and responsiveness to GA fluctuations. This adaptability allows DELLAs to interact with a wide array of proteins but also makes their functional study challenging: disruptions in DELLA function lead to pleiotropic effects across multiple pathways.

To address this complexity, understanding DELLA interactors provides valuable insights into DELLA’s nuanced functions and regulation. Building on this approach, we investigated novel OsSLR1 interactors, OsbHLH089 and OsbHLH094, members of the basic-helix-loop-helix transcription factor family. Our findings reveal that *OsbHLH089* and *OsbHLH094* are expressed in mature pollen grains, with the *Osbhlh089/94* double mutant displaying smaller, indehiscent anthers and non-viable pollen, indicating their redundant role in pollen development during its late stages.

Through ChIP-Seq and RNA-Seq analyses, we identified five target genes repressed by *OsbHLH089* and *OsbHLH094* (*OsTDL1A*, *OsSPS1*, *OsDGD2β*, *OspPGM*, and *OsDPE2*) that are essential for pollen and anther development. Interestingly, we also observed the binding of these TFs to the *OsSLR1* promoter. Notably, while other target genes were repressed, *OsSLR1* was induced, with a significant protein accumulation in the double mutant compared to the Kitaake background in the late stages of pollen development. This suggests that OsSLR1 accumulation may compromise pollen viability, further highlighting the critical regulatory role of *OsbHLH089* and *OsbHLH094* in repressing OsSLR1 levels for proper pollen and anther development.

## Introduction

DELLA proteins are key transcriptional regulators in plants, influencing processes from seed germination, plant height, and flowering to leaf senescence, fruit development, and stress responses. Their broad impact arises from their ability to interact with a wide array of transcriptional regulators and transcription factors (TFs). In Arabidopsis alone, the list of DELLA interacting partners already exceeds the 150 proteins (Marín-de La Rosa et al., 2014). For example, DELLA proteins form complexes with ABI3 and ABI5 to suppress germination (Lim et al., 2013), sequester PIF3 and PIF4 to limit hypocotyl elongation (De Lucas et al., 2008; Feng et al., 2008; Li et al., 2016a), delay flowering by forming complexes with FLOWERING LOCUS C (FLC) (Li et al., 2016b) and slow leaf senescence by binding WRKY45 (Chen et al. 2017). Additionally, DELLA enhances pathogen tolerance by interacting with JAZ repressors to release MYC2- dependent JA signaling (Kazan and Manners 2012).

Given the complexity of DELLA’s roles, characterizing specific DELLA interactors is key to understanding their full regulatory roles. In cereal crops, DELLAs accumulate in several stresses and play a role in yield; however, their pleiotropic effects limit the application of DELLA in crop improvement (Miralles et al., 1998a; Miralles et al., 1998b; Peng et al., 1999; de Vleesschauwer et al., 2016; Wang et al., 2020; Spielmeyer et al., 2022). Understanding DELLA function through their interacting partners could help harness DELLA functions to develop new strategies to improve crop productivity under climate change scenarios. Recently, we proposed two novel interactors of the single rice DELLA, OsSLR1: OsbHLH089 and OsbHLH094, members of the bHLH (basic Helix-Loop-Helix) transcription factor family, which is the second largest TF family in plants (Fernandes et al. 2024). In rice, 167 bHLH family members were identified and classified into 22 subfamilies (Li et al. 2006).

bHLH proteins possess a basic DNA-binding domain and a helix-loop-helix dimerization domain. The basic region enables binding to specific DNA sequences, while the helix-loop-helix motif facilitates dimerization, forming transactivation complexes that regulate gene expression. (Atchley et al. 1999). Typically, bHLH proteins bind to the E-box (5’-CANNTG-3’) sequence. A specific variant within the E-box family, the G-box (5’-CACGTG-3’), is recognized by approximately 81% of bHLH proteins (Atchley et al. 1999; Toledo-Ortiz et al. 2003; Li et al. 2006). bHLH TFs participate in many growth and development processes but with great representation in anther and pollen development (Nan et al. 2022; Ortolan et al. 2023).

In this study, we characterize the roles of OsbHLH089 and OsbHLH094 in rice pollen and anther development. Our findings indicate that these TFs act by modulating OsSLR1 levels, providing insights into DELLA’s regulatory network.

## Methods

### Plant material and growth conditions

The rice variety used in this study was *Oryza sativa* L. ssp. *Japonica* cv. Kitaake. Plants were grown in controlled-growth chamber conditions (12 h light at 28°C/12 h dark at 24°C, 400 μmol m^-2^ s^-1^) with a relative humidity of 60%.

*Osbhlh089* and *Osbhlh094* single and double mutants, as well as Kitaake plants, were vegetatively propagated from tillers.

### Transgene construct

*OsbHLH089* promoter region (2022 bp upstream the 5’UTR region) was subcloned into pHGWFS7. Overexpression constructs for OsbHLH089 and OsbHLH094 were assembled with a maize ubiquitin promoter and 3xFLAG tag into pHb7m34GW as the destination vector.

For *Osbhlh089* and *Osbhlh094* CRISPR/Cas9 mutants, sgRNAs targeting the first exon were designed using CRISPR-P (<u>crispr.hzau.edu.cn/CRISPR2/</u>) (Lei et al. 2014). DNA oligonucleotides were synthesized, annealed, and cloned into cut accepter vectors harboring the wheat U6 promoter, then assembled with Cas9 and gRNA vectors for transformation.

All primers used to generate transformation constructs and gRNAs are listed in Supplemental Tables 1 and 2, respectively.

The constructs were introduced into *Agrobacterium tumefaciens* EHA105 and underwent rice transformation.

### Rice transformation

Promoter-GUS reporter, gain, and loss-of-function lines were introduced into rice via Agrobacterium-mediated transformation of calli from scutellum tissue. Transformation followed the Langdale lab protocol described here (Carvalho et al. 2024). Transgenic lines with somaclonal variations were eliminated by successive selection through the T3 generation.

### Transformed lines evaluation

All lines were screened by the presence of Cas9 cassette by PCR to detect *hptII* gene. For *OsbHLH089* and *OsbHLH094* tagged lines, the levels of overexpressed protein were evaluated by Western Blot targeting the FLAG tag.

*OsbHLH089* and *OsbHLH094* single and double mutants were tested for the presence of insertions or deletions (InDels) in the target loci using PCR and Sanger sequencing (Brinkman et al. 2014). The chromatograms were analyzed using Synthego ICE software.

### Yeast two-hybrid

Yeast two-hybrid assays were performed as described in the manual of Matchmaker Gold Yeast Two-Hybrid Systems (Clontech). Full-length and several truncated forms of the OsbHLH089, OsbHLH094, and OsSLR1 were subcloned into pGBKT7 and pGADT7 vectors to construct different bait, respectively. The constructions were co-transformed into yeast strain Y2HGold (Clontech, US) by LiAc method (Gietz et al. 1995). Cells were grown on SD minimum, lacking leucine and tryptophan. To test possible interactions, transformed colonies were plated in SD, which lacked leucine, tryptophan, adenine, and histidine. To evaluate the ability of OsbHLH089 and OsbHLH094 to form homo and heterodimers, both CDS sequences were cloned into both Y2H vectors, and several combinations were transformed, as mentioned before. For the interaction screening, the transformed yeast colonies were plated in SD lacking leucine, tryptophan, and histidine and supplemented with 1 mM 3-amino-1,2,4-triazole (3’AT). The combinations with the empty vectors were used as negative controls. pAD-WT and pBD-WT from the HybriZAP 2.1 kit (Stratagene, USA) were used as controls of positive interaction in all Y2H assays.

### Subcellular localization

OsbHLH089 and OsbHLH094 were fused to GFP by subcloning into pH7FWG2,0. The constructs were transiently expressed in leaves of *Nicotiana benthamiana* strain by infiltration with *Agrobacterium tumefaciens* EHA105. The nuclear marker mCherry-NLS was used as a control.

Fluorescence signals were visualized using the Zeiss LSM 880 with Airyscan mode. Images were processed using Fiji (ImageJ) (Schindelin et al. 2012).

### Histochemical GUS staining

GUS activity was detected using the 5-bromo-4-chloro-3-indolyl-beta-D-glucuronic acid and cyclohexyl ammonium salt (X-Gluc) cleavage assay. Samples were incubated in 90% ice-cold acetone for 30 min at -20 °C, then incubated with staining solution (2mM X-Gluc, 100 mM phosphate buffer, 10 mM EDTA, 6 mM Ferrocyanide and 6 mM Ferricyanide). After staining, samples were fixed by incubating with ethanol: acetic acid (3:1) solution, followed by clearing solution (6M Urea, 30% Glycerol, and 0.1% Triton X-100). Images were obtained in a Leica DM 6000B optical microscope and an Axio Zoom V16 stereo microscope.

### RNA extraction and quantitative RT-PCR Analysis

Total RNA was extracted using the Direct-zol™ RNA MiniPrep (Zymo Research, Irvine, CA, USA) according to the manufacturer’s instructions. RNA from plant material with high starch content was performed using the Lithium Chloride (LiCl) precipitation method (Manickavelu et al. 2007). RNA integrity and genomic DNA contamination of all samples were examined by agarose gel electrophoresis. Quantification was done using a spectrophotometer (NanoDrop, ND-1000, Thermo Scientific, USA). The isolated total RNA was converted into complementary DNA (cDNA) using a first-strand cDNA synthesis kit (SuperScript III, Invitrogen). Reverse qRT-PCR was performed in a LightCycler system (Roche Molecular Biochemicals, Germany) using SYBR Premix Ex Taq (Takara, Japan). Relative expression levels were normalized against the expression levels of *UBC2, eF*-1a, and *EP* using the -2^ΔCt^ method. The qRT-PCR primers are listed in Supplementary Table 1.

### Phenotypic characterization of mutant plants

An A6000 Sony digital camera documented whole rice plant morphology. Spikelets, florets, and (in)dehiscent anthers were photographed with an Axio Zoom V16 stereo microscope. To get a clearer image of pollen content inside the anthers, freshly collected anthers were incubated for several days in a Clearing Solution (6 M Urea, 30% Glycerol, and 0.1% Triton X-100), and then images were taken using a Leica DM 6000B optical microscope. Anthers were tapped in a phosphate-buffered saline (PBS) solution to release pollen and assess pollen morphology. In the case of the *Osbhlh089/94* double mutant plants, the anther lodicules were gently opened, and pollen was released into the PBS solution. To obtain higher amounts of pollen, three anthers were needed, whereas in other genotypes, one was enough. Pollen viability was evaluated using the fluorescein diacetate (FDA) method developed by Li (Li 2011) under a Leica DM 6000B optical microscope. Around 350 pollen grains per 3 independent anthers from three independent plants (biological replicates) were counted at random (with or without fluorescence), evaluating a total of 1000 pollen grains per biological replicate. To calculate the percentage of pollen viability, the number of high fluorescence-emitting grains in each field was divided by the total amount of pollen grains assessed in the correspondent bright field image, and this quotient was multiplied by 100. To determine anther defects, we perform semi-thin sections of anthers during the late stages of development (stages 10-12). The stages were previously classified by Zhang and Wilson (Zhang and Wilson 2009). Briefly, spikelets were fixed in 3% (v/v) glutaraldehyde in 0.2 M sodium phosphate buffer (pH 7.0), then dehydrated in a graded series of ethanol solutions, embedded in Technovit 7100 resin (Hereaus Kulzer, Germany), polymerized at room temperature, and sectioned at 5 μm thickness using a microtome (Leica RM2235, Germany). The sections were stained with 0.05% (w/v) toluidine blue (Sigma) and photographed using a Leica DM 6000B optical microscope.

### Protein extraction and Western blot

For the Chip-Seq, total plant protein was extracted from 14-days-old plants of Kitaake, *OsbHLH089,* and OsbHLH94 tagged lines to evaluate overexpression and for OsSLR1 assessment from Kitaake and *Osbhlh089/94* double mutant total protein from anthers (stage 10-12) was collected.

Each sample was powdered in liquid nitrogen, and the total protein was extracted using the trichloroacetic acid (TCA)/acetone method (Niu et al., 2018).The protein content was quantified using the Pierce 660-nm Protein Assay Reagent (Thermo Scientific, USA) with bovine serum albumin as standard (Thermo Scientific, USA). To the tagged lines overexpression, fifteen micrograms were separated in a 12% SDS page gel. To evaluate OsSLR1 levels, ten micrograms of total protein were separated into a 10% SDS page gel. Both gels were transferred to PVDF Hybond-P membranes (Amersham, UK). The protein extracts from the tagged lines were probed with anti-FLAG monoclonal antibody (Sigma, #F3165; dilution 1:1 000) and sheep anti-mouse HRP conjugate antibody (NA 931 Amersham; 1:20 000). For SLR1 detection, a custom-made anti- SLR1 antibody (1:5 000) and anti-rabbit HRP conjugate antibody (A16096 Novex; 1:20 000) were used. To ensure that equal amounts of protein were loaded, the membranes were stripped by incubation with 2 M NaOH for 5 min and reprobed with anti-β-tubulin (sc-55529 Santa Cruz Biotechnology, 1:1 000) and sheep anti-mouse HRP conjugate antibody (NA 931 Amersham; 1:20 000).

The signals on blots were detected using ECL western blotting substrate (Amersham, UK).

### Multiple sequence alignment

Sequence amino acid alignment was performed by using CLUSTAL OMEGA online tool (https://www.ebi.ac.uk/Tools/msa/clustalo/) and visualized by the Jalview multiple alignment editor (Waterhouse et al. 2009).

### ChIP-Seq and data analysis

Three grams of leaf tissue from fully expanded leaves from 14 days of seedlings were fixed with 1% formaldehyde solution. Crosslinking was quenched by adding glycine solution to a final concentration of 0.125 M. Chromatin immunoprecipitation (ChIP) was performed as described (Gendrel et al. 2005), with the exception that chromatin-bound proteins were immunoprecipitated with Anti-FLAG® M2 magnetic beads (Sigma, USA) The ChIP-Seq libraries were prepared and sequenced by Illumina NovaSeq PE150 at Novogene (Cambridge, United Kingdom).

For data analysis, the reads were subjected to quality check in FastQC Toolkit. Adaptor and low-quality reads were further processed by trimmomatic software (Bolger et al. 2014), followed by the alignment to the *Oryza sativa* japonica reference genome (IRGSP-1.0) (Ensembl Plant v56) by Bowtie2 aligner (Langmead and Salzberg 2012). Peak calling was performed using MACS2 (Zhang et al., 2008). Bigwig files were used to visualize the peaks in the integrative genomics viewer. Annotation, visualization of ChIP peaks, and distribution of candidate binding regions across the rice genome were carried out using the R package ChIPseeker 1.36.0 version (Yu et al. 2015). Motif calling was performed using STREME with default parameters (Bailey, 2021).

### RNA-Seq and data analysis

Anthers were gathered and subjected to maceration using a tissue lyser. A portion of the resulting macerated anthers was observed under a light microscope to verify the effective shredding of pollen. Total RNAs were extracted with the Direct-zol™ RNA MiniPrep (Zymo Research, Irvine, CA, USA) from the late stages of anther development (stage 10-12) of Kitaake and *Osbhlh089/94* double mutant in three replicates. The stages were periodically validated through the utilization of semi-thin sections of anthers, as previously described, although the potential for some degree of intermingling remains present. RNA was quantified using Qubit™ RNA Assay Kits (Invitrogen) on Qubit® (Bio-Sciences, Dublin, Ireland), and RNA integrity was assessed using the Agilent 2100 Bioanalyzer (Agilent Technologies, USA). The RNA libraries were prepared, and samples were sequenced by Illumina NovaSeq PE150 at Novogene (Cambridge, United Kingdom). For data analysis, the reads were subjected to quality check in FastQC Toolkit. Adaptor and low-quality reads were further processed by trimmomatic software (Bolger et al. 2014), followed by genome annotation to the Oryza sativa japonica reference genome (IRGSP- 1.0) (Ensembl Plant v56) by STAR Feng et al., 2008; De Lucas et al., 2008. Differential expression analysis was performed using the DESeq2 R package. P-value < 0.01 and |log2 fold change| >2 was set as the threshold for significantly different expression in *Osbhlh089/94* mutant versus the Kitaake.

### Statistical analysis

The results of the study were expressed as mean±S.D. Data were analyzed by using a one-way analysis of variance test (ANOVA) followed by Dunnett’s t-test for multiple comparisons.

### Accession numbers

Sequence data from this article can be found in the Rice Genome Annotation Project Database under the following accession numbers: OsSLR1 (Os03g0707600), OsbHLH089 (Os03g0802900), OsbHLH094 (Os07g0193800), OsIDEF1 (Os08g0101000), UBC2 (Os03g0123100), eF-1a (Os03g0177500), and EP (Os05g0182700), OsTDL1A (Os12g0472500), OsSPS1 (Os01g0919400), OsDGD2β (Os03g0268300), OspPGM (Os10g0189100) and OsDPE2 (Os07g0662900).

## Results

### OsbHLH089 and OsbHLH094 are expressed in pollen grains

To investigate *OsbHLH089* expression patterns, we generated a *P-OsbHLH089* reporter construct with a 2.2 kb promoter region fused to the GUS reporter gene (Figure 1a). GUS staining revealed expression in mature pollen grains, developing seeds (1–5 DAA), and mature embryos DAA (Figure 1b, Supplementary Figure 1a). Due to technical limitations, we were unable to generate a *P-OsbHLH094* reporter construct. Therefore, to confirm *OsbHLH089* expression and assess *OsbHLH094* expression, we conducted qRT-PCR across various tissues, particularly focusing on late pollen development stages: stage 10 (vacuolated pollen stage), stage 11 (young bicellular pollen stage), and stage 12 (mature pollen stage (Zhang and Wilson, 2009). *OsbHLH089* showed peak expression at stage 12, early seed development (1, 3, and 5 DAA), and in mature embryos, with lower expression in other reproductive tissues. *OsbHLH094* also displayed elevated expression in anthers, followed by glumes and pistils, albeit with generally lower and more tissue-specific expression than *OsbHLH089* (Figure 1c).

**Figure 1.**
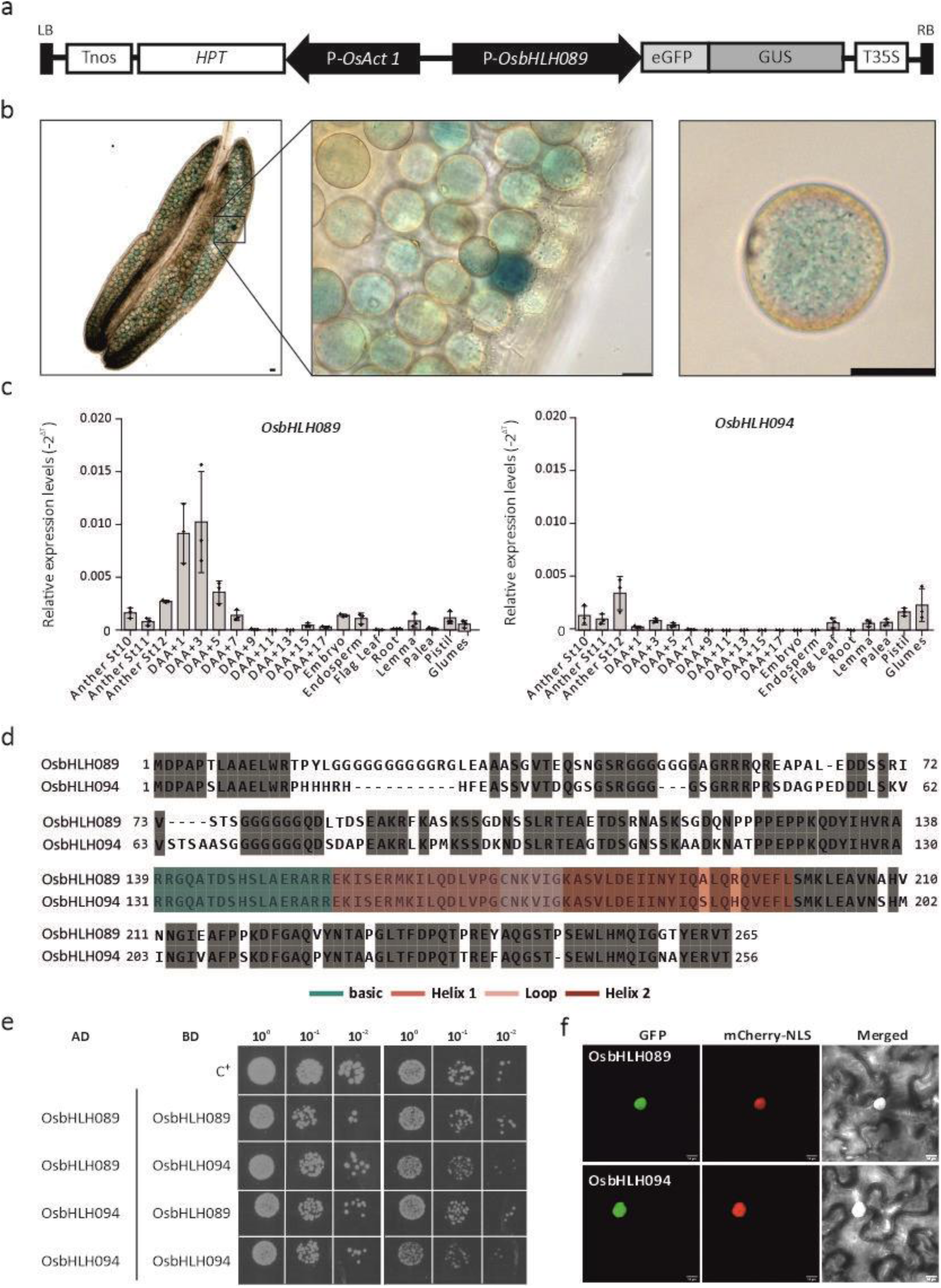
Gene expression profile of OsbHLH089 and OsbHLH094, dimerization and subcellular localization. (a) Schematic diagram of the P-OsbHLH089::GUS expression vector. (b) GUS histochemical staining in mature pollen grains. Spikelets were collected and stained. The anthers were removed to be photographed. Scale bar=25 µm. (c) qRT-PCR of OsbHLH089 (left) and OsbHLH094 (right) expression in various tissues of WT plants. Results are normalized against the expression levels of UBC2, eF-1a, and EP using the -2^ΔCt^ method. Individual values (dots) and means (bars) of three independent biological replicates are shown. (d) (d) protein sequence alignment of OsbHLH089 and OsbHLH094 was performed on CLUSTAL OMEGA online tool and visualized by the Jalview multiple alignment editor. Both sequences share 71% similarity. Green and three shades of pink boxes show the basic and HLH domains, respectively. (e) Y2H assay. Both OsbHLH089 and OsbHLH094 were fused with the GAL4 AD and with the GAL4 DNA BD. The appropriate vectors were transformed into the yeast Y2HGold strain. The different yeast strains were plated on a SD medium lacking Leu and Trp (SD/-Leu/-Trp) or on a SD medium lacking Trp, Leu, and His supplemented with 1 mM 3’AT (SD/-Leu/-Trp/-His) for the screening. pAD-WT/pBD- WT (Wild-type fragment C of lambda cI repressor) was used as a positive control (C+). As negative control both OsbHLH089 and OsbHLH094 were co-transformed with the suitable empty vector. The interaction was confirmed using three different clones. (f) Subcellular localization of OsbHLH089 and OsbHLH094 fused with GFP into appropriate expression vectors together with a vector harboring mcherry-NLS before Agrobacterium transfection of *N. benthamiana* leaves and analysis by using confocal laser scanning microscope. Scale bar=25 μm.

Evolutionary research on plant bHLH TFs has shown that OsbHLH089 and OsbHLH094 belong to the same clade (Catarino et al. 2016), indicating a high degree of similarity between the two proteins confirmed by protein alignment, especially in the bHLH domains (Figure 1d). Given the high similarity between the bHLH domains, we explored their dimerization potential. A Y2H assay showed that both TFs could form homo- and heterodimers (Figure 1e). To explore their subcellular localization, we expressed OsbHLH089 and OsbHLH094 fused with GFP in *N. benthamiana* leaves. *OsbHLH089* and *OsbHLH094* localize exclusively in the nucleus (Figure 1f).

### OsbHLH089 and OsbHLH094 regulate pollen fertility

To gain deeper insights into the function of OsbHLH089 and OsbHLH094, we generated single and double mutants using the CRISPR/Cas9 system (Supplementary Figure 2). *Osbhlh089* and *Osbhlh094* single mutants showed no effect on rice development, while the *Osbhlh089/94* double mutants produced no seeds (Figure 2 and Supplementary Figures 3 and 4). Morphological analysis of the florets of the *Osbhlh089/94* mutant plants revealed that the anthers were smaller and paler compared with the Kitaake and the single mutants (Figure 2a-c, Supplemental Figure 4a). The anthers also remained completely indehiscent without releasing mature pollen grains at anthesis, even after dissection (Figure 2d). Pollen was shrunken, without starch accumulation, and less abundant when compared to the other genotypes (Figure 2e). Moreover, the FDA viability staining showed that no viable pollen was generated in the double mutant (Figure 2f). In contrast, the single mutants did not exhibit any differences in terms of pollen viability, shape, or size compared to Kitaake pollen (Supplementary Figure 4e-f), suggesting that both OsbHLH089 and OsbHLH094 are redundant in their role in male gametophyte development.

**Figure 2.**
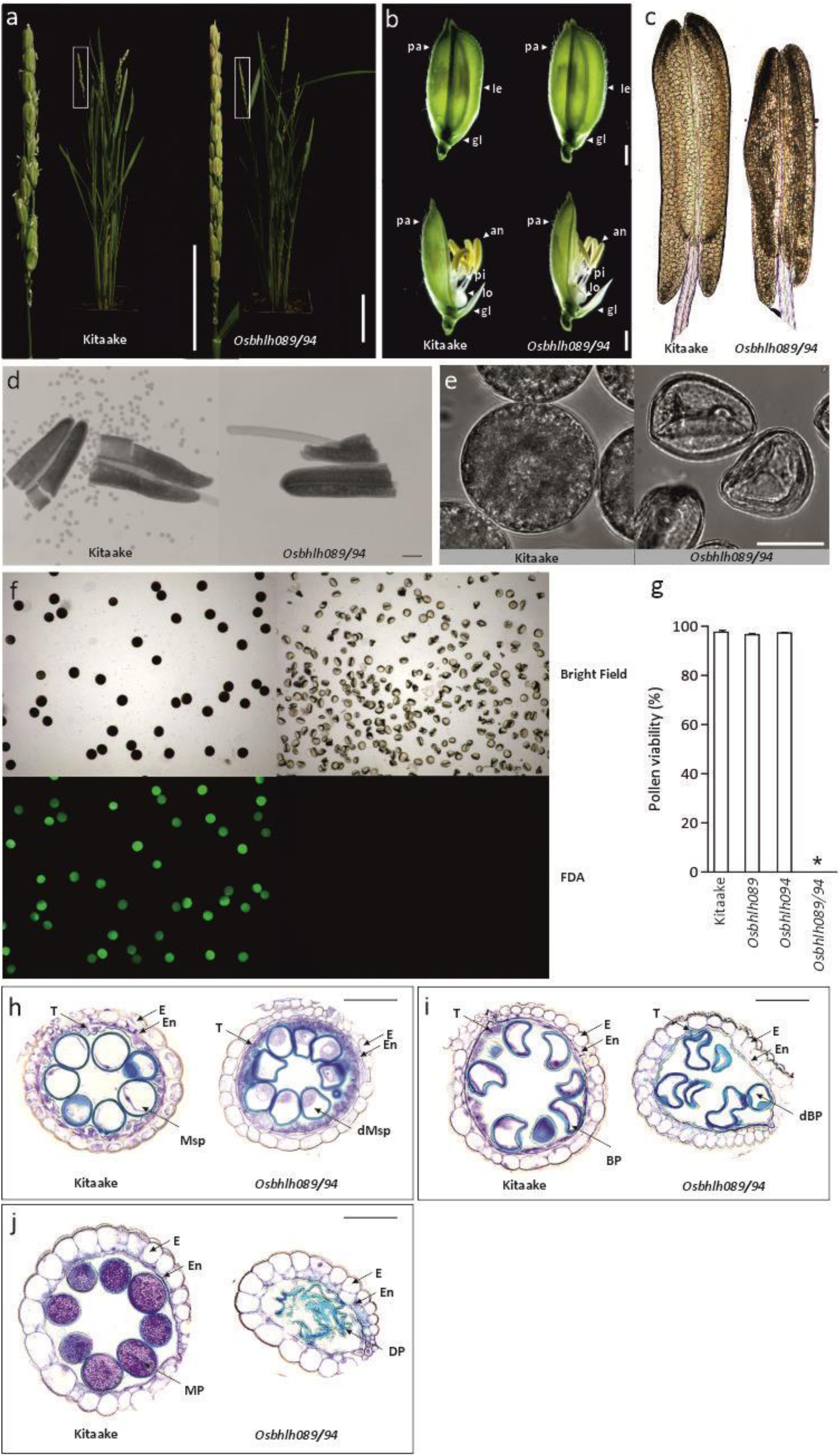
Phenotypic comparison between Kitaake and *Osbhlh089/94* mutant. (a) Kitaake and *Osbhlh089/94-1b* mutant plants after heading. On the left, a close-up of a representative panicle (white square). Scale bar=10 cm (plants) and scale bar=5 cm (panicle). (b) Spikelet (top) and floret (bottom) after heading. Scale bar=1 mm. (c) Anther morphology. Scale bar=0.1 mm. (d) Mature pollen after chopping the anthers. Scale bar=0.1 mm. (e) Mature pollen grains. Scale bar=25 µm. (f) Representative images of pollen viability test using fluorescein diacetate (FDA). Fluorescence indicates viable pollen. Scale bar=0.1 mm. (g) Quantification of pollen viability in Kitaake, single and double mutants pollen grains. (h-j) Locules from the semi-thin anther section of Kitaake and *Osbhlh089/94* from late stages of anther development. (h) Stage 10. (i) Stage 11. (j) Stage 10. BP, bicellular pollen; dBP, defective bicellular pollen dMsp, defective microspore: E, epidermis; En, endothecium; DP, degraded microspore; Msp, microspore; MP, mature pollen; T, tapetum. Scale bar=50 µm

To verify the impact of pollen defects observed in the *Osbhlh089/94* mutant on the development of the anthers, we performed semi-thin sections of anthers from stage 10 to stage 12 of anther development (Figure 4h-j). Semi-thin sections of anthers showed that in the double mutant, microspores were irregularly shaped, and tapetal cells were thicker at stage 10 compared with the Kitaake (Figure 4h). By stage 11, the anther tapetum layer disappeared prematurely (Figure 4i), and by stage 12, the locules contained collapsed pollen grains and an enlarged endothecium (Figure 4j).

These findings indicate that OsbHLH089 and OsbHLH094 are crucial for proper pollen and anther development.

### OsbHLH089 and OsbHLH094 bind to the promoters of anther/pollen-related genes

To identify OsbHLH089 and OsbHLH094 target genes, we performed ChIP-Seq experiments in stable rice lines overexpressing OsbHLH089 and OsbHLH094. The tagged lines showed no visible phenotypes. We identified that OsbHLH089 and OsbHLH094 binding sites are located mostly in promoter and distal intergenic regions (Figure 3a). The core motif with significant enrichment in the sequenced OsbHLH089 and OsbHLH094-binding regions was an E-box (CANNTG) (Figure 3b).

**Figure 3.**
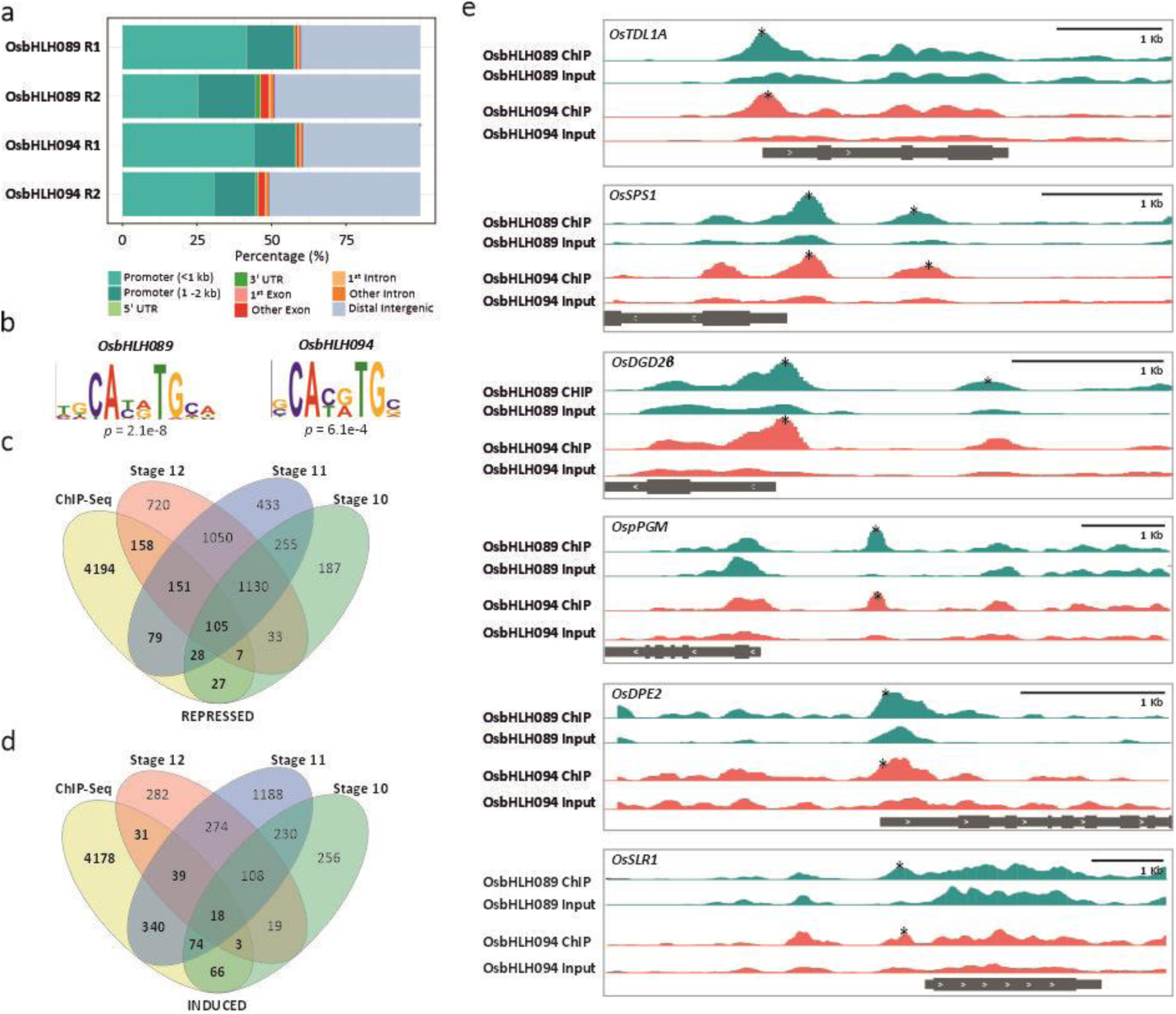
Identification of target genes of OsbHLH089 and OsbHLH094 by ChIP-Seq and RNA-Seq analysis. (a) Distribution of candidate OsbHLH089 and 94-binding regions across the rice genome. (b) Motif analysis using STREME to identify core enriched motifs. (c) Venn diagram showing the overlap between putative OsbHLH089 and OsbHLH094 target genes identified by ChIP-Seq and repressed genes identified in later stages of anther development of *Osbhlh089/94- 1b* double mutant relative to the Kitaake by RNA-Seq. (d) Venn diagram showing the overlap between putative OsbHLH089 and OsbHLH094 target genes identified by ChIP-Seq and induced genes identified in later stages of anther development of *Osbhlh089/94-1b* double mutant relative to the Kitaake by RNA-Seq. (e) Visualization of ChIP-Seq results of the putative OsbHLH089 and OsbHLH094 target genes repressed in late stages of anther development in *Osbhlh089/94-1b* double mutant. Genes, in grey, are shown below each panel, with boxes corresponding to exons and bars to UTR regions or introns. The asterisks represent the motif locations determined by MACS2.

**Figure 4.**
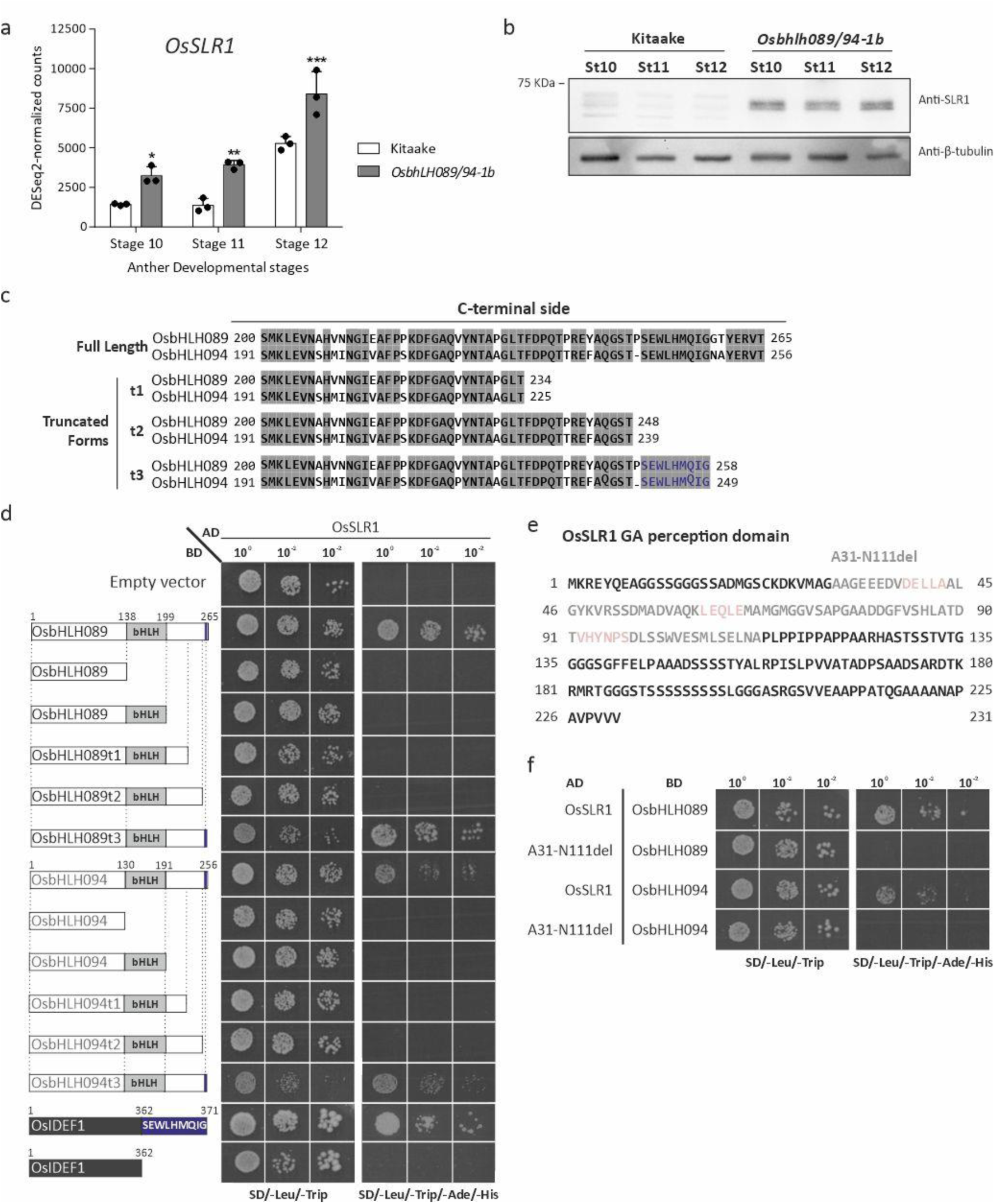
SLR1 accumulates in the *Osbhlh089/094* mutant. (a) OsSLR1 transcript levels in WT and *Osbhlh089/094* mutant on late stages of pollen development using DeSeq2-normalized counts. (b) Total protein extracts from anthers from stage 10-12 in WT and *Osbhlh089/94* double mutant immunoblotted with a custom-made anti-OsSLR1 antibody. Anti-β-tubulin was used as a loading control. (c) Schematic representation of the amino acid sequence alignment of three OsbHLH089 and OsbHLH094 C-terminal truncated forms used in this study. (d) Interaction between OsSLR1 and five OsbHLH089 and OsbHLH094 truncated forms: N-terminal side before the bHLH domain, N-terminal side including the bHLH domain, three C-terminal truncated forms (t1, t2, and t3) and OsIDEF1 and OsIDEF1 harboring the SEWLHMQIG motif. OsSLR1 against OsbHLH089 and OsbHLH094 full-length proteins was used as a positive control. (e) The amino acid sequence of OsSLR1 DELLA regulatory domain. Highlighted in pink are the motifs crucial for GID1 binding. The lighter amino acids were deleted to be tested in this study. (f) Interaction between the deleted version of OsSLR1 (A31-N111del) with OsbHLH089 and OsbHLH094 full-length proteins. OsSLR1 forms were fused with the GAL4 AD and OsbHLH089 and OsbHLH094 and derivatives with the GAL4 DNA BD into appropriate expression vectors before transfer into yeast (Y2HGold strain). The different yeast strains were plated on a SD medium lacking Leu and Trp (SD/-Leu/- Trp) or on a SD medium lacking Trp, Leu, Ade, and His (SD/-Leu/-Trp/-Ade/-His) for the screening. OsSLR1 against OsbHLH089 and OsbHLH094 full-length proteins was used as a positive control. BD alone was used as a negative control. The interaction was confirmed by three different clones.

Next, to narrow down the number of enriched peaks and select candidate target genes that could be relevant in anther development, we performed RNA-Seq analysis of three anther developmental stages (stage 10-12), comparing *Osbhlh089/94* mutant plants with Kitaake. (Supplementary Table 3). A principal component analysis based on transcriptomic profiles showed a clear separation of samples based on the line along PC1. Anther samples from Kitaake were clearly grouped according to the corresponding development stage, while samples from *Osbhlh089/94* mutant showed less divergence (Supplementary Figure 5a). Differential gene expression analysis between *Osbhlh089/94* and Kitaake lines in each developmental stage indicated that Stage 11 is the condition with a higher number of DEGs, followed by stage 12 and stage 10 (Supplementary Figure 5b).

Afterward, we compare the DEGs identified as downregulated (repressed) and upregulated (induced) by the double knockout and genes with promoter regions (2000 bp upstream transcription start site) overlapping the common OsbHLH089/094-enriched peaks from the ChIP-Seq analysis (**Error! Reference source not found.**e 6c-d). In the Venn diagram with the repressed DEGs, we observed an overlap of 422 DEGs at stage 12 with the ChIP-Seq experiment, 363 with stage 11 and 164 with stage 10 (Figure 3c, Supplementary Table 4). Looking at the induced DEGs, we observed an overlap of 91 DEGs between stage 12 with OsbHLH089 and OsbHLH094 enriched peaks, 531 with stage 11, and 161 with stage 10. (Figure 3d, Supplementary Table 5). Among the 4364 repressed DEGs, we detected five potential OsbHLH089 and OsbHLH094 targets genes for anther and pollen development: *tapetum determinant 1 (OsTDL1A), sucrose phosphate synthase enzyme 1 (OsSPS1), digalactosyldiacylglycerol synthase (OsDGD2β), plastidic phosphoglucomutase (OspPGM),* and *isproportionating enzyme2 (OsDPE2)*. *OsTDL1A* is specific to stage 10 of pollen development, *OsSPS1* to stage 11, and *OsDGD2β* to stage 12. *OspPGM* is expressed during stages 10 and 11, while *OsDPE2* is expressed between stages 11 and 12.

From a total of 2928 induced DEGs, and based on the predicted functional annotation, we found no direct association with anther or pollen development.

### OsbHLH089 and OsbHLH094 regulate OsSLR1 levels

To investigate the potential transcriptional regulation of OsSLR1 by OsbHLH089 and OsbHLH094, we analyzed ChIP-Seq data for enrichment in the SLR1 promoter region. Interestingly, our findings reveal that OsbHLH089 and OsbHLH094 can bind directly to the OsSLR1 promoter (Figure 3e). OsSLR1 was identified as one of the DEGs in our RNA-Seq dataset, with fold change values ranging from 0.65 to 1.50, depending on the stage. Normalization of read counts showed that OsSLR1 expression was upregulated in the *Osbhlh089/94* mutant compared to Kitaake across all stages of pollen development (Figure 4a). Since the DELLA proteins are known to be regulated by ubiquitin-mediated proteolysis, transcript and protein levels might not always correlate. Therefore, we assessed the protein levels in the three stages of pollen development and showed that while no OsSLR1 protein was detected in any of the stages from the Kitaake, there was a massive accumulation in all the stages in the double knockout (Figure 4b). These results suggest that OsbHLH089 and OsbHLH094 play a role in the regulation of OsSLR1.

DELLA proteins bind to bHLH domains, blocking bHLH TFs from targeting gene promoters (De Lucas et al. 2008; Feng et al. 2008a). To explore in more detail the interaction between OsbHLH089 and OsbHLH094 with OsSLR1, we conducted a Y2H assay (Figure 4, Supplemental Figure 6). *OsSLR1* was fused to GAL4 AD and five truncated TF forms were fused to GAL4 BD (Figure 4c-d). *OsSLR1* interacted with the C-terminal truncated form of *OsbHLH089* and *OsbHLH094,* containing the SEWLHMQIG motif. This motif alone, fused to *OsIDEF1*, a TF that was previously shown not to interact with OsSLR1 (Fernandes et al. 2024), was sufficient for interaction with *OsSLR1* (Figure 4c).

The GRAS domain in DELLA proteins is known to mediate protein-protein interactions (Marín-de La Rosa et al., 2014). To assess whether this applies in our case, we deleted critical motifs within the DELLA regulatory domain, specifically the DELLA, LEQLE, and VHYNPS sequences (Figure 4e). This deletion notably disrupted the interaction between OsbHLH089/OsbHLH094 and OsSLR1 (Figure 4f), indicating that the integrity of the DELLA regulatory domain is crucial for binding.

## Discussion

DELLA proteins exert their function by partnering with transcriptional regulators or TFs, so understanding the role of DELLA interacting partners is key to shed light on DELLAs’ myriad functions and complex regulation. Here, we performed the functional characterization of OsbHLH089 and OsbHLH094, previously identified novel OsSLR1 interactors (Fernandes et al. 2024), and demonstrated their important role in pollen and anther development.

Our findings establish that *OsbHLH089* and *OsbHLH094* are expressed in mature pollen grains. Although we did not observe high expression levels during earlier stages of pollen development, we detected early pollen defects as soon as stage 10 (vacuolated pollen stage) and stage 11 (young bicellular pollen stage). The presence of shrunken, non-viable pollen grains in the *Osbhlh089/94* double mutant anthers underscores the essential role these TFs play in rice male reproductive structures. The absence of notable phenotypic differences between the *OsbHLH089* and *OsbHLH094* single mutants compared to the *Osbhlh089/94* double mutant indicates functional redundancy in male reproductive development. This redundancy may explain the low expression levels of these TFs since reduced expression could help maintain the stable functional redundancy observed in these duplicated genes (Qian et al. 2010).

To further elucidate the molecular functions of OsbHLH089 and OsbHLH094, we identified genome-wide binding sites of these TFs *in vivo* using ChIP-Seq, complemented with RNA-Seq analysis. Our results revealed 1024 potential target genes for OsbHLH089 and OsbHLH094. Among these, we highlighted genes with well-established roles in pollen and anther development, including *OsTDL1A, OsSPS1, OsDGD2β, OspPGM*, and *OsDPE2*. These transcripts were significantly repressed in the *Osbhlh089/94* double mutant, suggesting that OsbHLH089 and OsbHLH094 are required for the activation of the transcription of these genes to problably ensure proper pollen and anther development. On the other hand, we also found that OsbHLH089 and OsbHLH094 bind to the promoter of *OsSLR1*. In the double mutant, OsSLR1 transcript levels were upregulated throughout all stages of pollen development, accompanied by a substantial accumulation of OsSLR1 protein.

Several studies highlight the critical role of GA metabolism in anthers and pollen development. Mutants with impaired GA biosynthesis consistently show underdeveloped anthers and non-viable pollen (Koornneef and van der Veen 1980; Nester and Zeevaart 1988; Jacobsen and Olszewski 1991; Goto and Pharis 1999; Murray et al. 2003; Kaneko et al. 2004). Loss-of-function of *OsCPS1, GID1*, and *GID2* results in DELLA protein accumulation, suggesting that this accumulation disrupts tapetum and pollen development (Aya et al. 2009) .

Our findings suggest two possible scenarios for the observed non-viable pollen phenotype. First, *OsbHLH089* and *OsbHLH094* may be necessary for activating genes critical for anther and pollen development. Second, without *OsbHLH089* and *OsbHLH094*, *OsSLR1* is no longer repressed, leading to *OsSLR1* accumulation and the disruption of pollen development, possibly by sequestering TFs required to activate pollen-related genes showcasing a that. *OsbHLH089* and *OsbHLH094* may act as both activators and repressors, as many TFs exhibit dual roles (Boyle and Després 2010). However, it is possible that *OsbHLH089* and *OsbHLH094* may not independently target the *OsSLR1* promoter. This hypothesis arises from our observations of tagged lines used for ChIP-Seq analysis: when *OsbHLH089* and *OsbHLH094* are driven by a constitutive promoter, the plants show no distinct phenotype. If *OsbHLH089* and *OsbHLH094* were directly repressing *OsSLR1* alone, their expression across various tissues and developmental stages would likely result in a dwarf phenotype. Given our Y2H results, showing that *OsbHLH089* and *OsbHLH094* can form homo- or heterodimers, we aim to assess whether their regulatory action occurs as dimers. However, it is evident that heterodimer formation is not essential for the function of these TFs. If heterodimerization were required, we would expect to see a pollen-impaired phenotype in the single mutants, similar to what is observed in the double mutant.

Additionally, we explored the interaction of OsSLR1 with OsbHLH089 and OsbHLH094 by Y2H using truncated forms of these TFs. DELLA proteins are known to bind HLH domains, blocking the binding of bHLH TFs to target promoters (De Lucas et al., 2008; Feng et al., 2008). However, our results revealed that the HLH domain is not responsible for interacting with OsSLR1. While OsSLR1 does not directly interfere with the DNA-binding domain, it may influence DNA binding via conformational changes at the TFs’ C-terminal regions. We also found that the SEWLHMQIG motif that is conserved in both TFs is sufficient for the interaction between OsSLR1 with OsbHLH089 and OsbHLH094. Identifying motifs responsible for DELLA interactions could allow *in silico* predictions of new DELLA interacting proteins. Next, we showed that DELLA motifs that are responsible for the interaction with GA-GID1 (DELLA, LEQLE, and VHYNPS) are essential in the interaction between OsSLR1 with OsbHLH089 and OsbHLH094. GRAS domain was always considered the DELLA domain that promotes interactions, and this ability has been conserved since DELLAs appearance in early land plants throughout evolution (Briones-Moreno et al. 2023) .The fact that the DELLA domain seems to play a role in this novel interaction with the bHLHs raised the question: Could the recruitment of DELLA to the GA-GID1-DELLA complex also have facilitated the establishment of novel interactions with other partners? It is clear that the GRAS domain is the main driver of DELLAs interactions; however, most DELLA interacting proteins have been identified through Y2H screenings where part of the DELLA domain was deleted because of DELLAs’ transactivation properties (Hou et al. 2010; Marín-de La Rosa et al., 2014); Marín-de la Rosa et al. 2015). This fact probably excluded the identification of N-terminal binding DELLA partners. Thus, it may be necessary to revise the strategies for DELLA-interacting partner identification, ensuring that full-length DELLAs are tested.

Our research reinforces the importance of DELLA proteins in male reproduction, particularly in later stages of anther development. We observed that DELLA accumulation during these stages disrupts viable pollen production and causes male infertility, supporting previous findings. Conversely, early-stage pollen development shows the opposite effect: *OsSLR1* loss-of-function mutants exhibit sterility due to premature tapetum cell death, leading to microspore degradation by stage 10 and lack of mature pollen (Ikeda et al. 2001, Jin et al. 2022).Our work clarifies that *OsSLR1* has a stage-dependent role in anther and pollen development, partnering with *OsMS188* to activate pollen wall formation genes in early stages (Jin et al. 2022), while unregulated *OsSLR1* levels in later stages impair pollen viability by sequestering TFs required for gene activation related to pollen and anther development.

In conclusion, our findings underscore the finely-tuned balance between GA and DELLA proteins required for anther and pollen development, regulated by temporal expression. Our data suggest that in the later stages, DELLA proteins may inhibit male reproduction, highlighting the need for precise regulation of DELLA activity to maintain fertility and produce viable pollen.

## Author contributions

TF participated in the conception of research, executed experiments, analised the data and wrote the article. PMB participated in the bioinformatics analysis of ChIP-seq and RNA-seq data. MFT participated in the microscopy analysis and image adquisition of the reproductive analysis. PC guided execution of experiments. HS supported in the histologial preparations. JB provided suggestions on analysis, organization and writing. IAA conceived the original project, supervised experiments, and finalized the article. All authors discussed the results and commented on the manuscript.

### Acknowledgments

This work was supported by FCT - Fundação para a Ciência e a Tecnologia, I.P., through: GREEN- IT Bioresources for Sustainability R&D Unit base (DOI: 10.54499/UIDB/04551/2020) and programmatic (DOI: 10.54499/UIDP/04551/2020) funding; LS4FUTURE Associated Laboratory (DOI: 10.54499/LA/P/0087/2020); PhD fellowship awarded to TF (PD/BD/135584/2018) and PC (PD/BD/128403/2017), the Post-Doc contract awarded to PMB (DOI: 10.54499/DL57/2016/CP1369/CT0029), contract awarded to MFT though the European Union’s HORIZON-MSCA-2021-PF-01-01 programme [grant 101059247]. HS was supported through the FCT project PTDC/ASP-PLA/2007/2020. The funding sources had no involvement in study design, analyses, and interpretation of data, writing, or in the submission decision.

## Supplementary figures

**Supplementary Figure 1.**
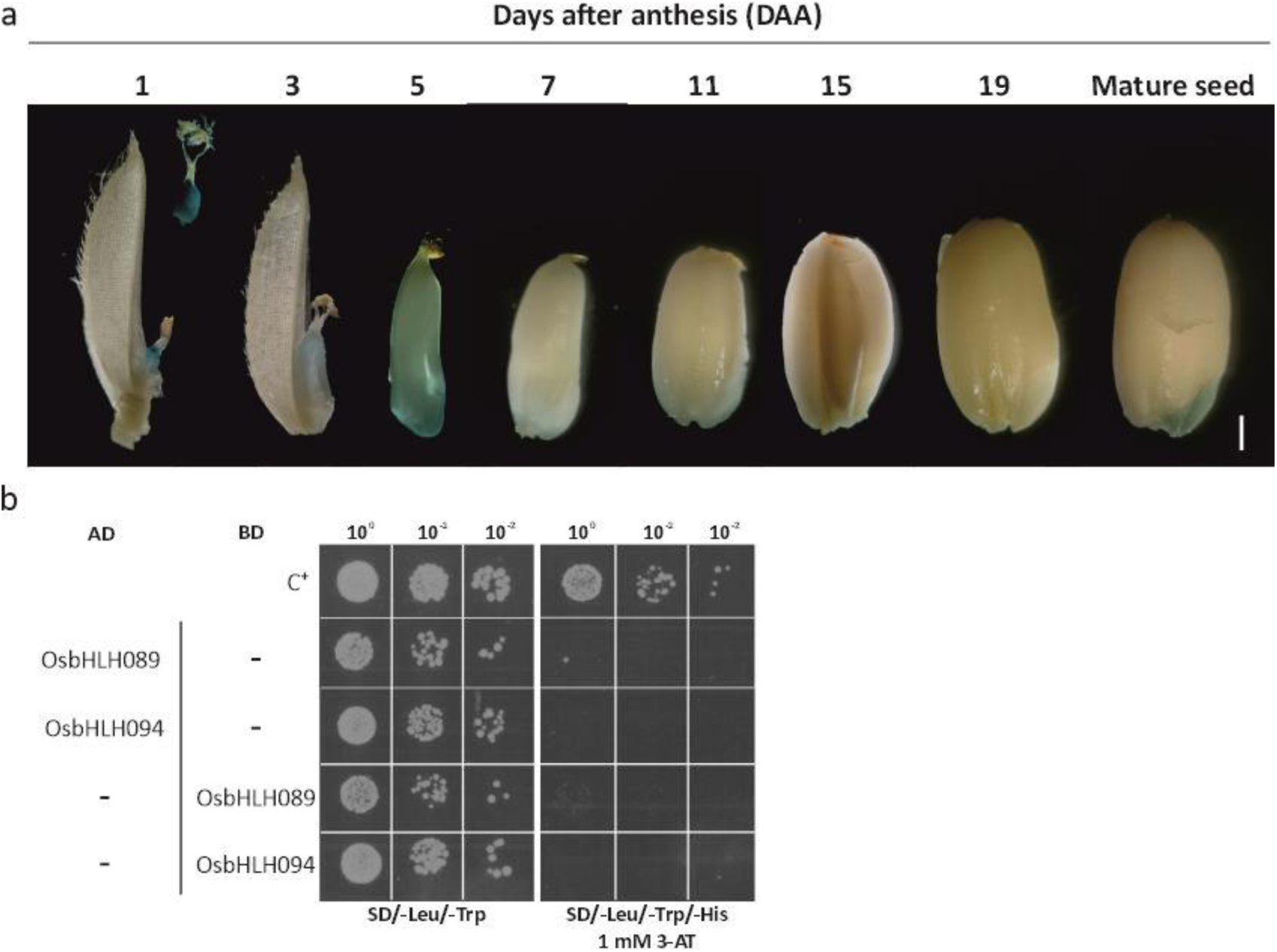
Gene expression profile of OsbHLH089:β-glucuronidase (GUS) reporter line. (a) GUS expression patterns during seed development. Rice spikelets were collected after anthesis (DAA) every two days. After staining, the lemma and palea were partially or completely removed from each spikelet during seed development. The stained spikelets were preserved in 70% ethanol and photographed. Scale bar=1 mm. (b) Y2H assay negative controls for OsbHLH089 and OsbHLH094 dimerization. Both OsbHLH089 and OsbHLH094 were fused with the GAL4 AD and with the GAL4 DNA BD. The appropriate vectors were transformed into the yeast Y2HGold strain. The different yeast strains were plated on a SD medium lacking Leu and Trp (SD/-Leu/-Trp) or on a SD medium lacking Trp, Leu, and His supplemented with 1 mM 3’AT (SD/-Leu/-Trp/-His) for the screening. pAD-WT/pBD-WT (Wild-type fragment C of lambda cI repressor) was used as a positive control (C+). As negative control both OsbHLH089 and OsbHLH094 were co-transformed with the suitable empty vector. The interaction was confirmed using three different clones.

**Supplementary Figure 2.** CRISPR/Cas9-induced OsbhLH089 and OsbHLH094 single and double mutants. (a) Schematic illustration of CRISPR/Cas9 DNA constructs used in this study. *The Hygromycin B phosphotransferase* (*hptII*) gene, driven by Cauliflower Mosaic Virus 35 promoter (P-CaMV35S) was used as a selection marker. *Streptococcus pyogenes*-derived *Cas9* together with a nuclear localization signal (SpCAS9-NLS) and two sgRNAs: sgRNA1 targeting *OsbHLH089* and sgRNA2 targeting *OsbHLH094*. SpCAS9-NLS and gRNA expression were driven by the maize ubiquitin promoter (P-ZmUbi) and wheat U6 promoter (P-TaU6), respectively. Tnos, Nopaline Synthase terminator; T35S, Cauliflower Mosaic Virus 35S terminator. (b) Schematic diagram of the mutation details of *OsbHLH089* (top) and *OsbHLH094* (bottom) single mutant lines. (c) Schematic diagram of the mutation details of *OsbHLH089* (top) and *OsbHLH094* (bottom) double mutant line. (b-c) Schematic map of the genomic region of OsbHLH089 and OsbHLH094 and the sgRNA target site. Black boxes show exons, black lines show introns and white boxes show 5′- and 3′-untranslated regions (UTR). The grey box shows a hypothetical 5’UTR region, and the green region represents the bHLH domain. The red arrow indicates the CRISPR/Cas9 target site, and the black arrows show the position of PCR primers used for mutation detection. Sequence alignment of the sgRNA target region showing altered bases in different OsbHLH089 mutant lines compared with the WT. Dashed lines represent base deletions and bold nucleotides indicate base insertions. Red bases indicate the sgRNA targeting sequence and the underlined black bases indicate Protospacer Adjacent Motif (PAM) sites. The sequence alignment near the target site of *OsbHLH089* and *OsbHLH094* in the mutants and WT lines is also shown.

**Supplementary Figure 3.**
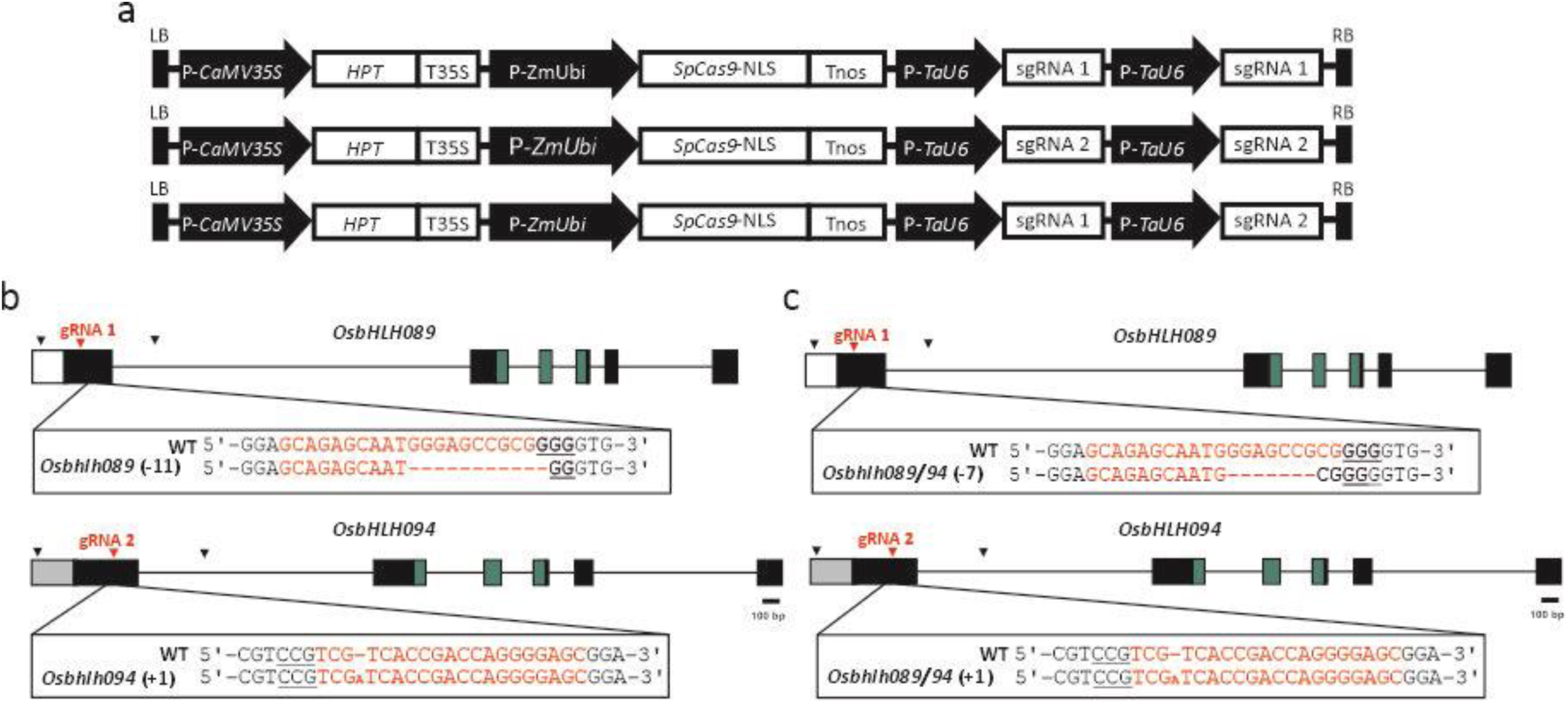
*Osbhlh089* and *Osbhlh094* single mutants have no impact on plant height or seed development. (a) The phenotype of *Osbhlh089* and *Osbhlh094* single mutants and Kitaake 60 DAG (days after germination). Scale bar=10 cm. (b) Plant height during the vegetative stage. (c) Flowering time is measured in days. (d) % filled seeds per total of seeds. (e) Panicle phenotype. Scale bar=5 cm. (f) Panicle length. (g) Seed phenotype. Scale bar=1 cm. (h) seed length. (i) 1000-grain weight. (j) seed width. Data are presented as means ± S.D. Box plots in (c,d,f, and i) show median (horizontal line) and individual values (black dots) (n=12 biological replicates). Data in (h and j) are presented as means ± S.D. (n=100 biological replicates). Comparisons between the single mutants and Kitaake were carried out by one-way analysis of variance (ANOVA) followed by Dunnett’s t-test.

**Supplementary Figure 4.**
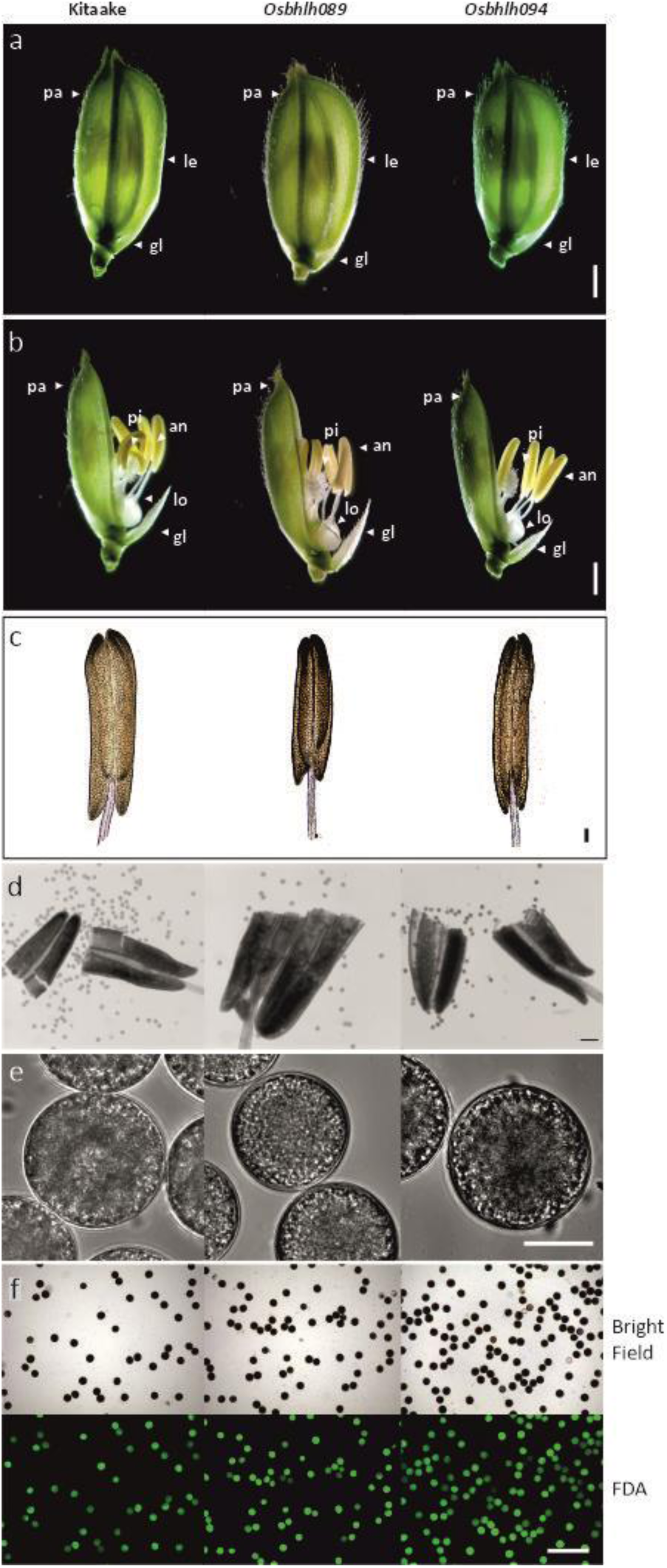
Phenotypic comparison between Kitaake, *Osbhlh089,* and *Osbhlh094* single mutants. (a) Spikelet morphology after heading. (b) Floret morphology after heading. Scale bar=1 mm. (c) Anther morphology. Scale bar=0.1 mm. (d) Mature pollen after chopping the anthers. Scale bar=0.1 mm. (e) Mature pollen structure from confocal laser scanning microscope. Scale bar=25 µm. (f) Representative images of pollen viability test using fluorescein diacetate (FDA). Fluorescence indicates viable pollen, and the absence of fluorescence shows dead pollen. Scale bar=0.1 mm.

**Supplementary Figure 5.**
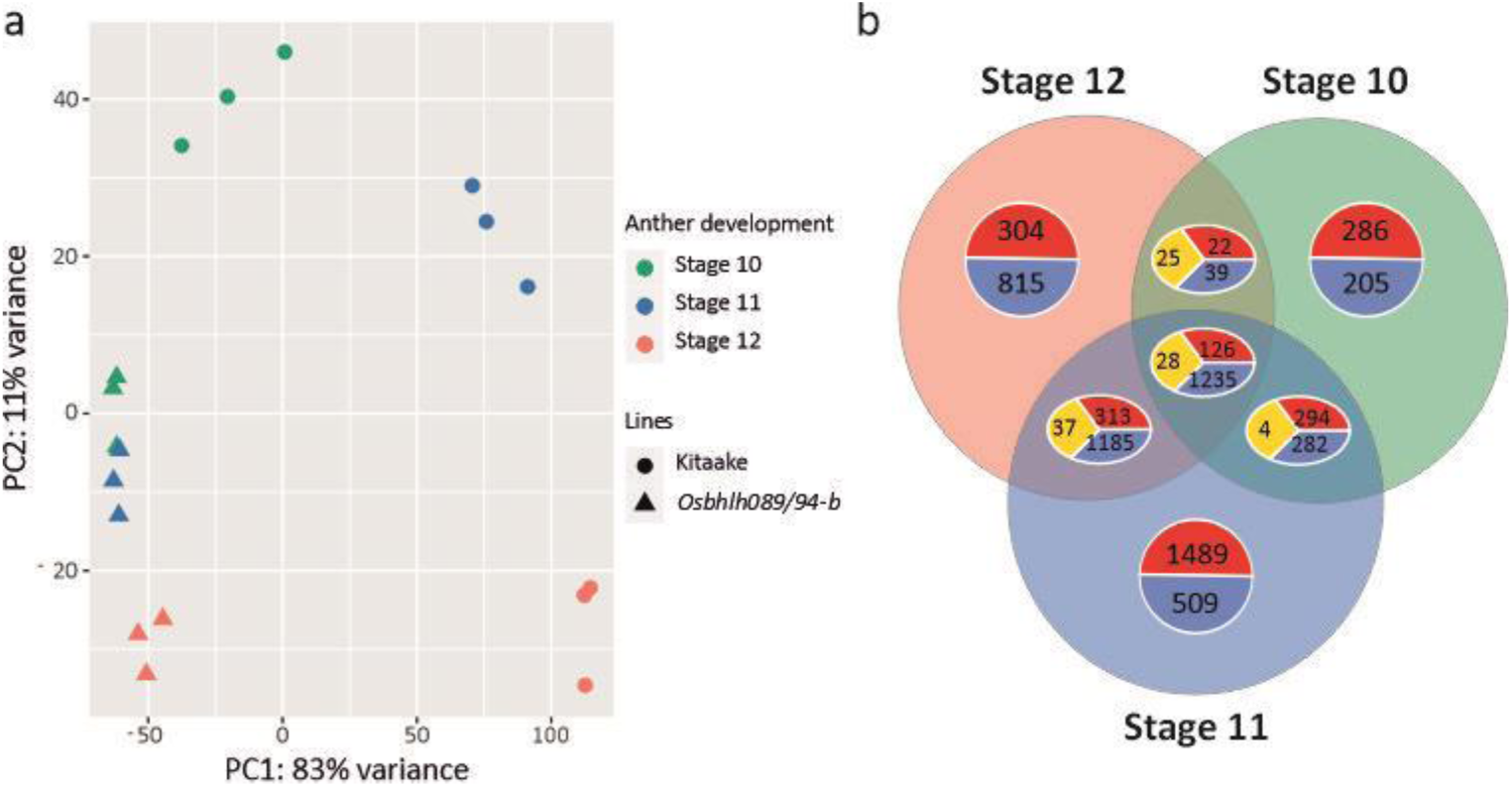
RNA-Seq of late stages of anther development in Kitaake and *Osbhlh089/94* mutant. (a) Principal Components analysis (PCA) of RNAseq data among different anther developmental stages (stage10-12) in Kitaake and *Osbhlh089/94* mutant. (b) Venn diagram showing DEGs in the *Osbhlh089/94* mutant. Foldchange > 2, p-value < 0.01. Red: upregulated genes; Blue: downregulated genes; Yellow: contrasting gene expression.

**Supplementary Figure 6.**
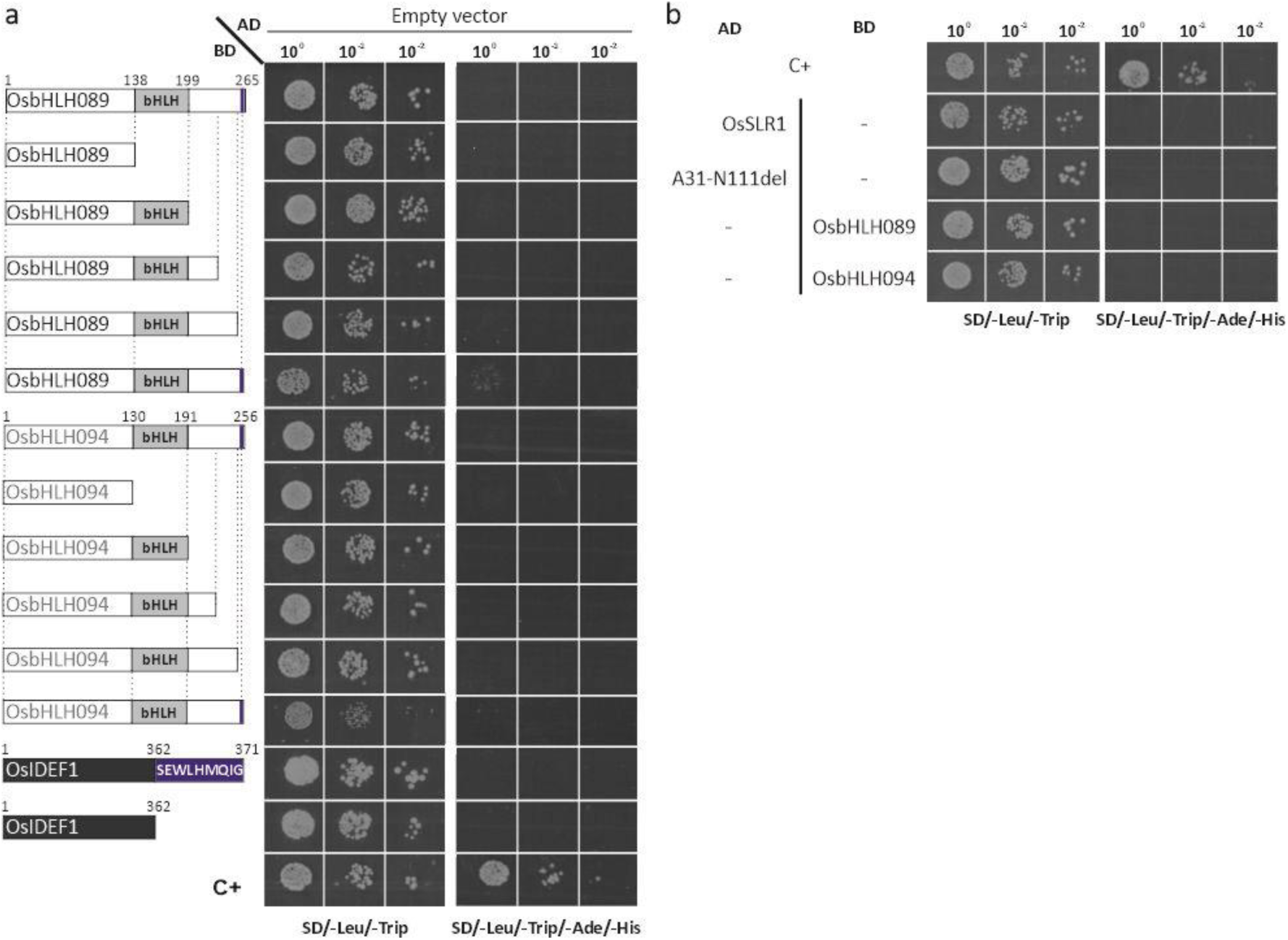
Protein-protein interaction negative controls for Y2H assays. (a) AD not interacting with OsbHLH089 and OsbHLH094 truncated forms: N-terminal side before the bHLH domain, N-terminal side including the bHLH domain, three C-terminal truncated forms (t1, t2, and t3) and OsIDEF1 and OsIDEF1 harboring the SEWLHMQIG motif. The different yeast strains were plated on a SD medium lacking Leu and Trp (SD/-Leu/-Trp) or on a SD medium lacking Trp, Leu, Ade, and His (SD/-Leu/-Trp/-Ade/-His) for the screening. pAD-WT/pBD-WT (Wild-type fragment C of lambda cI repressor) was used as a positive control (C+). The result was confirmed by three different clones. (b) AD not interacting with OsbHLH089 and OsbHLH094 and the binding domain (BD) is not interacting with OsSLR1 and the OsSLR1A31-N111del version. The different yeast strains were plated on a SD medium lacking Leu and Trp (SD/-Leu/-Trp) or on an SD medium lacking Trp, Leu, Ade, and His (SD/-Leu/-Trp/-Ade/-His) for the screening. pAD- WT/pBD-WT (Wild-type fragment C of lambda cI repressor) was used as a positive control (C+). The result was confirmed by three different clones.

## References

1. Atchley WR, Terhalle W, and Dress A. Positional Dependence, Cliques, and Predictive Motifs in the bHLH Protein Domain. J Mol Evol. 1999:48(5):501–516. 10.1007/PL00006494

2. Aya K, Ueguchi-Tanaka M, Kondo M, Hamada K, Yano K, Nishimura M, and Matsuoka M. Gibberellin Modulates Anther Development in Rice via the Transcriptional Regulation of GAMYB. Plant Cell. 2009:21(5):1453–1472. 10.1105/tpc.108.062935

3. Bailey TL. STREME: accurate and versatile sequence motif discovery. Bioinformatics. 2021:37(18). 10.1093/bioinformatics/btab203

4. Bolger AM, Lohse M, and Usadel B. Trimmomatic: a flexible trimmer for Illumina sequence data. Bioinformatics. 2014:30(15):2114–2120. 10.1093/bioinformatics/btu170

5. Boyle P and Després C. Dual-function transcription factors and their entourage. Plant Signal Behav. 2010:5(6):629–634. 10.4161/psb.5.6.11570

6. Brinkman EK, Chen T, Amendola M, and van Steensel B. Easy quantitative assessment of genome editing by sequence trace decomposition. Nucleic Acids Res. 2014:42(22):e168–e168. 10.1093/nar/gku936

7. Briones-Moreno A, Hernández-García J, Vargas-Chávez C, Blanco-Touriñán N, Phokas A, Úrbez C, Cerdán PD, Coates JC, Alabadí D, and Blázquez MA. DELLA functions evolved by rewiring of associated transcriptional networks. Nat Plants. 2023. 10.1038/s41477-023-01372-6

8. Carvalho P, Gomes C, Gonçalves I, Lourenço TF, Vlad D, Langdale JA, and Saibo NJM. The bHLH transcription factor OsPRI1 activates the *Setaria viridis PEPC1* promoter in rice. New Phytologist. 2024:241(6):2495–2505. 10.1111/nph.19556

9. Catarino B, Hetherington AJ, Emms DM, Kelly S, and Dolan L. The Stepwise Increase in the Number of Transcription Factor Families in the Precambrian Predated the Diversification of Plants on Land. Mol Biol Evol. 2016:33(11):2815–2819. 10.1093/molbev/msw155

10. Chen L, Xiang S, Chen Y, Li D, and Yu D. Arabidopsis WRKY45 Interacts with the DELLA Protein RGL1 to Positively Regulate Age-Triggered Leaf Senescence. Mol Plant. 2017:10(9):1174–1189. 10.1016/j.molp.2017.07.008

11. Feng S, MArtinez C, Gusmaroli G, Wang Y, Zhou J, Wang F, Chen L, Yu L, and Deng WX. Coordinated regulation of Arabidopsis thaliana development by light and gibberellins. Nature. 2008a:23(1):1–7. 10.1038/jid.2014.371

12. Feng S, Martinez C, Gusmaroli G, Wang Y, Zhou J, Wang F, Chen L, Yu L, Iglesias-Pedraz JM, Kircher S, et al. Coordinated regulation of Arabidopsis thaliana development by light and gibberellins. Nature. 2008b:451(7177):475–479. 10.1038/nature06448

13. Fernandes T, Gonçalves NM, Matiolli CC, Rodrigues MAA, Barros PM, Oliveira MM, and Abreu IA. SUMOylation of rice DELLA SLR1 modulates transcriptional responses and improves yield under salt stress. Planta. 2024:260(6):136. 10.1007/s00425-024-04565-1

14. Gendrel A-V, Lippman Z, Martienssen R, and Colot V. Profiling histone modification patterns in plants using genomic tiling microarrays. Nat Methods. 2005:2(3):213–218. 10.1038/nmeth0305-213

15. Gietz RD, Schiestl RH, Willems AR, and Woods RA. Studies on the transformation of intact yeast cells by the LiAc/SS-DNA/PEG procedure. Yeast. 1995:11(4):355–360. 10.1002/yea.320110408

16. Goto N and Pharis RP. Role of gibberellins in the development of floral organs of the gibberellin-deficient mutant, ga1-1, of *Arabidopsis thaliana*. Canadian Journal of Botany. 1999:77(7):944–954. 10.1139/b99-090

17. Hou X, Lee LYC, Xia K, Yan Y, and Yu H. DELLAs Modulate Jasmonate Signaling via Competitive Binding to JAZs. Dev Cell. 2010:19(6):884–894. 10.1016/j.devcel.2010.10.024

18. Ikeda A, Ueguchi-Tanaka M, Sonoda Y, Kitano H, Koshioka M, Futsuhara Y, Matsuoka M, and Yamaguchi J. Slender rice, a constitutive gibberellin response mutant, is caused by a null mutation of the SLR1 gene, an ortholog of the height-regulating gene GAI/RGA/RHT/D8. Plant Cell. 2001:13(5):999–1010. 10.1105/tpc.13.5.999

19. Jacobsen SE and Olszewski NE. Characterization of the Arrest in Anther Development Associated with Gibberellin Deficiency of the *gib-1* Mutant of Tomato. Plant Physiol. 1991:97(1):409–414. 10.1104/pp.97.1.409

20. Kaneko M, Inukai Y, Ueguchi-Tanaka M, Itoh H, Izawa T, Kobayashi Y, Hattori T, Miyao A, Hirochika H, Ashikari M, et al. Loss-of-Function Mutations of the Rice *GAMYB* Gene Impair α-Amylase Expression in Aleurone and Flower Development. Plant Cell. 2004:16(1):33–44. 10.1105/tpc.017327

21. Kazan K and Manners JM. JAZ repressors and the orchestration of phytohormone crosstalk. Trends Plant Sci. 2012:17(1):22–31. 10.1016/j.tplants.2011.10.006

22. Koornneef M and van der Veen JH. Induction and analysis of gibberellin sensitive mutants in Arabidopsis thaliana (L.) heynh. Theoretical and Applied Genetics. 1980:58(6):257–263. 10.1007/BF00265176

23. Langmead B and Salzberg SL. Fast gapped-read alignment with Bowtie 2. Nat Methods. 2012:9(4). 10.1038/nmeth.1923

24. Lei Y, Lu L, Liu HY, Li S, Xing F, and Chen LL. CRISPR-P: A web tool for synthetic single-guide RNA design of CRISPR-system in plants. Mol Plant. 2014:7(9):1494–1496. 10.1093/mp/ssu044

25. Li K, Yu R, Fan LM, Wei N, Chen H, and Deng XW. DELLA-mediated PIF degradation contributes to coordination of light and gibberellin signalling in Arabidopsis. Nat Commun. 2016a:7. 10.1038/ncomms11868

26. Li M, An F, Li W, Ma M, Feng Y, Zhang X, and Guo H. DELLA proteins interact with FLC to repress flowering transition. J Integr Plant Biol. 2016b:58(7):642–655. 10.1111/jipb.12451

27. Li X. Pollen Fertility/viability Assay Using FDA Staining. Bio. 2011:101(e75).

28. Li X, Duan X, Jiang H, Sun Y, Tang Y, Yuan Z, Guo J, Liang W, Chen L, Yin J, et al. Genome-Wide Analysis of Basic/Helix-Loop-Helix Transcription Factor Family in Rice and Arabidopsis. Plant Physiol. 2006:141(4):1167–1184. 10.1104/pp.106.080580

29. Lim S, Park J, Lee N, Jeong J, Toh S, Watanabe A, Kim J, Kang H, Kim DH, Kawakami N, et al. ABA- insensitive3, ABA-insensitive5, and DELLAs interact to activate the expression of SOMNUS and other high-temperature-inducible genes in imbibed seeds in Arabidopsis. Plant Cell. 2013:25(12):4863– 4878. 10.1105/tpc.113.118604

30. De Lucas M, Davière JM, Rodríguez-Falcón M, Pontin M, Iglesias-Pedraz JM, Lorrain S, Fankhauser C, Blázquez MA, Titarenko E, and Prat S. A molecular framework for light and gibberellin control of cell elongation. Nature. 2008:451(7177):480–484. 10.1038/nature06520

31. Manickavelu A, Kambara K, Mishina K, and Koba T. An efficient method for purifying high quality RNA from wheat pistils. Colloids Surf B Biointerfaces. 2007:54(2):254–258. 10.1016/j.colsurfb.2006.10.024

32. Marín-de la Rosa N, Pfeiffer A, Hill K, Locascio A, Bhalerao RP, Miskolczi P, Grønlund AL, Wanchoo-Kohli A, Thomas SG, Bennett MJ, et al. Genome Wide Binding Site Analysis Reveals Transcriptional Coactivation of Cytokinin-Responsive Genes by DELLA Proteins. PLoS Genet. 2015:11(7). 10.1371/journal.pgen.1005337

33. Marín-de La Rosa N, Sotillo B, Miskolczi P, Gibbs DJ, Vicente J, Carbonero P, Oñate-Sánchez L, Holdsworth MJ, Bhalerao R, Alabadí D, et al. Large-scale identification of gibberellin-related transcription factors defines group VII ETHYLENE RESPONSE FACTORS as functional DELLA partners. Plant Physiol. 2014a:166(2):1022–1032. 10.1104/pp.114.244723

34. Marín-de La Rosa N, Sotillo B, Miskolczi P, Gibbs DJ, Vicente J, Carbonero P, Oñate-Sánchez L, Holdsworth MJ, Bhalerao R, Alabadí D, et al. Large-scale identification of gibberellin-related transcription factors defines group VII ETHYLENE RESPONSE FACTORS as functional DELLA partners. Plant Physiol. 2014b:166(2):1022–1032. 10.1104/pp.114.244723

35. Miralles DJ, Katz SD, Colloca A, and Slafer GA. Floret development in near isogenic wheat lines differing in plant height. Field Crops Res. 1998:59:20–31.

36. Miralles DJ, Calderini DF, Pomar KP, and D’ambrogio2 A. Dwarfing genes and cell dimensions in different organs of wheat.

37. Murray F, Kalla R, Jacobsen J, and Gubler F. A role for HvGAMYB in anther development. The Plant Journal. 2003:33(3):481–491. 10.1046/j.1365-313X.2003.01641.x

38. Nan G-L, Teng C, Fernandes J, O’Connor L, Meyers BC, and Walbot V. A cascade of bHLH-regulated pathways programs maize anther development. Plant Cell. 2022:34(4):1207–1225. 10.1093/plcell/koac007

39. Nester JE and Zeevaart JAD. Flower development in normal tomato and a gibberellin-deficient (ga-2) mutant. Am J Bot. 1988:75(1):45–55. 10.1002/j.1537-2197.1988.tb12160.x

40. Niu L, Zhang H, Wu Z, Wang Y, Liu H, Wu X, and Wang W. Modified TCA/acetone precipitation of plant proteins for proteomic analysis. PLoS One. 2018:13(12):e0202238. 10.1371/journal.pone.0202238

41. Ortolan F, Trenz TS, Delaix CL, Lazzarotto F, and Margis-Pinheiro M. bHLH-regulated routes in anther development in rice and Arabidopsis. Genet Mol Biol. 2023:46(3 suppl 1). 10.1590/1678-4685-gmb-2023-0171

42. Peng J, Richards DE, Hartley NM, Murphy GP, Devos KM, Flintham JE, Beales J, Fish LJ, Worland AJ, Pelica F, et al. “Green revolution” genes encode mutant gibberellin response modulators. Nature. 1999:400(6741):256–261. 10.1038/22307

43. Qian W, Liao B-Y, Chang AY-F, and Zhang J. Maintenance of duplicate genes and their functional redundancy by reduced expression. Trends in Genetics. 2010:26(10):425–430. 10.1016/j.tig.2010.07.002

44. Schindelin J, Arganda-Carreras I, Frise E, Kaynig V, Longair M, Pietzsch T, Preibisch S, Rueden C, Saalfeld S, Schmid B, et al. Fiji: An open-source platform for biological-image analysis. Nat Methods. 2012:9(7):676–682. 10.1038/nmeth.2019

45. Spielmeyer W, Ellis MH, and Chandler PM. Semidwarf (sd-1), “‘green revolution’” rice, contains a defective gibberellin 20-oxidase gene. PNAS. 2022:99(13):9043–9048.

46. Toledo-Ortiz G, Huq E, and Quail PH. The Arabidopsis Basic/Helix-Loop-Helix Transcription Factor Family[W]. Plant Cell. 2003:15(8):1749–1770. 10.1105/tpc.013839

47. Vleesschauwer D, Seifi HS, Filipe O, Haeck A, Huu SN, Demeestere K, and Höfte M. The DELLA protein SLR1 integrates and amplifies salicylic acid- and jasmonic acid-dependent innate immunity in rice. Plant Physiol. 2016:170(3):1831–1847. 10.1104/pp.15.01515

48. Wang J, Qin H, Zhou S, Wei P, Zhang H, Zhou Y, Miao Y, and Huang R. The ubiquitin-binding protein OsDSK2a mediates seedling growth and salt responses by regulating gibberellin metabolism in rice. Plant Cell. 2020:32(2):414–428. 10.1105/tpc.19.00593

49. Waterhouse AM, Procter JB, Martin DMA, Clamp M, and Barton GJ. Jalview Version 2—a multiple sequence alignment editor and analysis workbench. Bioinformatics. 2009:25(9):1189–1191. 10.1093/bioinformatics/btp033

50. Yu G, Wang L-G, and He Q-Y. ChIPseeker: an R/Bioconductor package for ChIP peak annotation, comparison and visualization. Bioinformatics. 2015:31(14). 10.1093/bioinformatics/btv145

51. Zhang DB and Wilson ZA. Stamen specification and anther development in rice. Chinese Science Bulletin. 2009:54(14):2342–2353. 10.1007/s11434-009-0348-3

52. Zhang Y, Liu T, Meyer CA, Eeckhoute J, Johnson DS, Bernstein BE, Nusbaum C, Myers RM, Brown M, Li W, et al. Model-based Analysis of ChIP-Seq (MACS). Genome Biol. 2008:9(9). 10.1186/gb-2008-9-9-r137

