## Supplemental Tables for "OsbHLH089 and OsbHLH094 Modulate OsSLR1 Levels to Maintain Male Reproductive Fitness in Rice"

**Supplementary Tables**

**Supplementary Table 1** List of primers used.

| Primer Name | Direction | Sequence (5’-3’) | Notes |
| --- | --- | --- | --- |
| BHLH089-FW-GW | Forward | GGGGACAAGTTTGTACAAAAAAGCAGGCTTAATGGACCCCGCCCCCACGCT | Gateway cloning |
| BHLH089-RV-GW | Reverse | GGGGAC CAC TTT GTA CAA GAAAGCTGGGTTTCATGTCACTCTTTCATAGG | Gateway cloning |
| BHLH089_GW_ST_Rv | Reverse | GGGGACCACTTTGTACAAGAAAGCTGGGTCTGTCACTCTTTCATAGGTGC | Gateway cloning |
| BHLH094_synt_ST-GW-Fw | Forward | GGGGACAAGTTTGTACAAAAAAGCAGGCTTAATGGACCCTGCACCGAGTTT | Gateway cloning |
| BHLH094_synt_ST-GW-Rv | Reverse | GGGGACCACTTTGTACAAGAAAGCTGGGTCAGTAACCCTCTCATAGGCGTT | Gateway cloning |
| GTW_OsSLR1_FW | Forward | GGGGACAAGTTTGTACAAAAAAGCAGGCTCGATGAAGCGCGAGTACCAAGA | Gateway cloning |
| GTW_OsSLR1_Rv | Reverse | GGGGACCACTTTGTACAAGAAAGCTGGGTCCTAAACTTTGCGATCACGCCGC | Gateway cloning |
| bHLH089_gRNA_Fw | Forward | TACTAGTGTACAGTGCCACT | Plant genotyping |
| bhLH089_gRNA_Rv | Reverse | CCAGAACTACAGATTGGCTC | Plant genotyping |
| bHLH089_gRNA_seq_fw2 | Forward | TCGCCTCAAACCCTAGAGCT | Plant genotyping+sequencing |
| bHLH094_gRNA_fw2 | Forward | CATCACAGCTCTTCTTCTCC | Plant genotyping |
| bhLH094_gRNA_Rv | Reverse | GACACTGGCATGGACTTGGC | Plant genotyping |
| bHLH094_gRNA_Fw | Forward | TCCGCCTCGCCGCCACAT | Plant genotyping+sequencing |
| hptII_Fw | Forward | AATAGCTGCGCCGATGGTTTCTACA | *hptII* amplification |
| hptII_Rv | Reverse | AACATCGCCTCGCTCCAGTCAATG | *hptII* amplification |
| IDEF-FW-GW | Forward | GGGGACAAGTTTGTACAAAAAAGCAGGCTTAATGGGGCAAATGGACGGCGG | Gateway cloning |
| IDEF-RV-GW | Reverse | GGGGACCACTTTGTACAAGAAAGCTGGGTTCTAGGGATTTGTTGTCTGC | Gateway cloning |
| IDEF_SEWLHMQIG_Gw_rv | Reverse | GGGGACCACTTTGTACAAGAAAGCTGGGTC CTAACCAATCTGCATGTGGAGCCATTCCGAGGGATTTGTTGTCTGC | Gateway cloning |
| bHLH089_17truncated_muta_fw | Forward | GCCATTCCGATCATGGTGTTGATCCCTGGGC | Mutagenesis |
| bHLH089_17truncated_muta_rv | Reverse | GCCCAGGGATCAACACCATGATCGGAATGGC | Mutagenesis |
| bHLH094_17truncated_muta_fw | Forward | GCCACTCCGATCAAGTACTCCCTTGCGCAAAC | Mutagenesis |
| bHLH094_17truncated_muta_rv | Reverse | GTTTGCGCAAGGGAGTACTTGATCGGAGTGGC | Mutagenesis |
| bHLH089_8truncated_muta_fw | Forward | GGCTCCACATGCAGATTGGTTGAGGCACCTATGAA | Mutagenesis |
| bHLH089_8truncated_muta_rv | Reverse | TTCATAGGTGCCTCAACCAATCTGCATGTGGAGCC | Mutagenesis |
| bHLH094_8truncated_muta_fw | Forward | CCCTCTCATAGGCGTTCAGCCTATTTGCATGTGG | Mutagenesis |
| bHLH094_8truncated_muta_rv | Reverse | CCACATGCAAATAGGCTGAACGCCTATGAGAGGG | Mutagenesis |
| bHLH089-N-GW_RV | Reverse | GGGGACCACTTTGTACAAGAAAGCTGGGTTCATGCCCTCACATGGATGTAGTC | Gateway cloning |
| bHLH089-domain-RV | Reverse | GGGGACCACTTTGTACAAGAAAGCTGGGTTCACAAAAACTCAACTTGCCGTTGC | Gateway cloning |
| bHLH094-N-GW-synt-Rv | Reverse | GGGGACCACTTTGTACAAGAAAGCTGGGTTCAGGCCCGCACATGGATATAATC | Gateway cloning |
| bHLH089_trun1_G694T_fw | Forward | GTGGATCAAACGTCAATCAAGGAGCCGTGTTGTATAC | Mutagenesis |
| bHLH089_trun1_G694T_rv | Reverse | GTATACAACACGGCTCCTTGATTGACGTTTGATCCAC | Mutagenesis |
| bHLH94_N+domain_del574-577_fw | Forward | CCGCCTCCAGTTTCATAAGAACTCGACTTGATGC | Mutagenesis |
| bHLH94_N+domain_del574-577_rv | Reverse | GCATCAAGTCGAGTTCTTATGAAACTGGAGGCGG | Mutagenesis |
| bHLH94_trun1_del673_fw | Forward | GGGTCGAAAGTCAGCCGGCCGCTGTG | Mutagenesis |
| bHLH94_trun1_del673_rv | Reverse | CACAGCGGCCGGCTGACTTTCGACCC | Mutagenesis |
| bHLH089_qPCR_fw | Forward | ACATCCATGTGAGGGCAAGAAGG | qRT-PCR |
| bHLH089_qPCR_rv | Reverse | TTTCTCACGACGTGCCCTTTCG | qRT-PCR |
| bHLH094_qPCR_fw | Forward | GCACGGCGAGAGAAGATAAGTG | qRT-PCR |
| bHLH094_qPCR_rv | Reverse | GACCTTATTGCATCCAGGGACAAG | qRT-PCR |
| OsEF-1a fw | Forward | TGGTGACCAAGATCGACAGA | qRT-PCR |
| OsEF-1a rv | Reverse | GCATCACCGTTCTTGAGGA | qRT-PCR |
| OsEP_qPCRa_fw | Forward | TGAGCAAAATGGTGGAAAGC | qRT-PCR |
| OsEP_qPCRa_rv | Reverse | CAGTTGCAACCCCTGTATGA | qRT-PCR |
| UBC2-qF | Forward | TTGCATTCTCTATTCCTGAGCA | qRT-PCR |
| UBC2-qR | Reverse | CAGGCAAATCTCACCTGTCTT | qRT-PCR |

**Supplementary Table 2** List of guide RNAs (gRNAs) used for CRISPR/Cas9 plant genome engineering. gRNA targets and oligonucleotide sequences of gRNA target sites (SpCas9) and PAM sequence (5'-NGG-3') are underlined.

| gRNA name | Targeted gene | Sequence (5’-3’) + PAM |
| --- | --- | --- |
| gRNA 1 | *OsbHLH089* | GCAGAGCAATGGGAGCCGCG GGG |
| gRNA 2 | *OsbHLH094* | GCTCCCCTGGTCGGTGACGA CGG |

**Supplementary Table 3** Significantly different expression in the *Osbhlh089/94* mutant compared to Kitaake, identified using the DESeq2 R package with a P-value threshold of < 0.01

**[Please see Supplementary Table 3.xlsx file]**

**Supplementary Table 4** Putative *OsbHLH089/94* target genes repressed in the *OsbHLH089/94* mutant**.**

**[Please see Supplementary Table 4.xlsx file]**

**Supplementary Table 5** Putative *OsbHLH089/94* target genes assigned induced in the *OsbHLH089/94-1b* mutant.

**[Please see Supplementary Table 5.xlsx file]**
